# Disentangling interactions among mercury, immunity, and infection in a Neotropical bat community

**DOI:** 10.1101/2020.06.12.135475

**Authors:** Daniel J. Becker, Kelly A. Speer, Jennifer M. Korstian, Dmitriy V. Volokhov, Hannah F. Droke, Alexis M. Brown, Catherene L. Baijnauth, Ticha Padgett-Stewart, Hugh G. Broders, Raina K. Plowright, Thomas R. Rainwater, M. Brock Fenton, Nancy B. Simmons, Matthew M. Chumchal

## Abstract

1. Contaminants such as mercury are pervasive and can have immunosuppressive effects on wildlife. Impaired immunity could be important for forecasting pathogen spillover risks, as many land-use changes that generate mercury contamination also bring wildlife into close contact with humans and domestic animals. However, the interactions among contaminants, immunity, and infection are difficult to study in natural systems, and empirical tests of possible directional relationships remain rare.
2. We capitalized on extreme mercury variation in a diverse bat community in Belize to test association among contaminants, immunity, and infection. By comparing a previous dataset of bats sampled in 2014 with new data from 2017, representing a period of rapid agricultural land conversion, we first confirmed bat species more reliant on aquatic prey had higher fur mercury. Bats in the agricultural habitat also had higher mercury in recent years. We then tested covariation between mercury and cellular immunity and determined if such relationships mediated associations between mercury and common bacterial pathogens. As bat ecology can dictate exposure to mercury and pathogens, we also assessed species-specific patterns in mercury–infection relationships.
3. Across the bat community, individuals with higher mercury had fewer neutrophils but not lymphocytes, suggesting stronger associations with innate immunity. However, the odds of infection for hemoplasmas and *Bartonella* spp. were generally lowest in bats with high mercury, and relationships between mercury and immunity did not mediate infection patterns. Mercury also showed species- and clade-specific relationships with infection, being associated with especially low odds for hemoplasmas in *Pteronotus mesoamericanus* and *Dermanura phaeotis.* For *Bartonella* spp., mercury was associated with particularly low odds in the genus *Pteronotus* but high odds in the Stenodermatinae.
4. *Synthesis and application:* Lower general infection risk in bats with high mercury despite weaker innate defense suggests contaminant-driven loss of pathogen habitat (i.e., anemia) or vector mortality as possible causes. Greater attention to these potential pathways could help disentangle relationships among contaminants, immunity, and infection in anthropogenic habitats and help forecast disease risks. Our results also suggest contaminants may increase infection risk in some taxa but not others, emphasizing the importance of considering surveillance and management at different phylogenetic scales.

## Introduction

Wildlife are commonly exposed to many contaminants that are ubiquitous in the environment, including heavy metals, organic compounds, and pesticides (Smith et al., 2007). Contaminants can be novel stressors that have direct impacts such as mortality (Davidson, 2004) and more subtle consequences such as immunosuppression (Grasman, 2002). For example, even relatively low concentrations of mercury (Hg), a neurotoxic heavy metal, can impair wildlife immunity (Scheuhammer, Meyer, Sandheinrich, & Murray, 2007). When immunosuppression manifests in increased susceptibility to pathogens, environmental gradients of contaminants could increase infection prevalence or intensity in wild hosts (Becker, Albery, et al., 2020). These elevated infection risks could be especially relevant in the context of environmental changes such as gold mining and agricultural land clearing, both of which are associated with bioaccumulation of contaminants like Hg in aquatic and terrestrial food webs (Farella, Davidson, Lucotte, & Daigle, 2007; Palheta & Taylor, 1995). As land conversions such as these also facilitate novel interactions among wildlife, domestic animals, and humans, impaired immunity could further increase pathogen spillover risks (Borremans, Faust, Manlove, Sokolow, & Lloyd-Smith, 2019).

The interactions among contaminants, immunity, and infection are notoriously difficult to study in natural systems, and empirical tests of possible directional relationships remain rare (Ross, 2002). Without captive or field experiments, natural systems must demonstrate sufficient variation in contaminant exposure and infection status to permit inference, and these criteria may be hard to meet in practice. For example, recent work on urbanized bobcats demonstrated strong effects of anticoagulant exposure on immunity in ways that should increase susceptibility, but pathogens of clinical relevance were generally rare, making it difficult to draw epidemiological inferences (Serieys et al., 2018). Additionally, effects may be difficult to detect if contaminant concentrations have low variance or are below toxicity thresholds (Fisk et al., 2005). Other examples suggest contaminants may instead decrease infection risks (Prüter et al., 2018), but the degree to which such patterns might be mediated by contaminant effects on immunity are unclear. Field-based assessments of how variable contaminant concentrations are associated with immunity and common pathogens are necessary to disentangle directional relationships.

Here, we capitalized on high variation in Hg and infection across species in a diverse Neotropical bat community in Belize to test associations among contaminants, immunity, and infection. Hg concentrations are typically highest in aquatic animals because methylmercury (MeHg), the bioaccumulative form of Hg, is produced in aquatic ecosystems (Chumchal et al., 2011). However, such contaminants can transfer into terrestrial ecosystems when terrestrial consumers feed on aquatic prey contaminated with MeHg (Cristol et al., 2008). Neotropical bats are ecologically diverse (Gunnell & Simmons, 2012; Rojas, Vale, Ferrero, & Navarro, 2011), and their diet variation enables strong heterogeneity in Hg exposure. Specifically, our past work showed that how often species feed on aquatic prey (or prey with some life stages linked to aquatic ecosystems) determines bat fur Hg, such that insectivores and species feeding on amphibians and fish have greater diet exposure than frugivores and sanguivores (Becker, Chumchal, et al., 2018). Such variation should produce strong associations with immunity, as even sublethal Hg concentrations correlate with immunity (Becker et al., 2017). Because bats can be vulnerable to extracellular pathogens and can also harbor viral and bacterial zoonoses (Brook & Dobson, 2015), immune differences driven by dietary variation in Hg could have implications for disease risks to and from bats. Lastly, this region in Belize is also undergoing intensive land clearance for agriculture similar to much of Latin America (Patterson, 2016), and thus analyses of Hg, immunity, and infection could help assess how land use affects wildlife and human health.

Here we built upon our prior studies of Hg and of infectious disease in Neotropical bats (e.g., Becker, Chumchal, et al., 2018; Becker, Speer, et al., 2020) to address three study aims. First, we combined historic and new data on Hg concentrations in bat fur, an indication of long-term metal exposure (Flache et al., 2015), and compared contaminant load over a three-year period and two sites. This greater within-species sample size allowed us to more robustly assess whether bat dietary connectivity to aquatic ecosystems predicts Hg bioaccumulation and if such patterns persist despite spatiotemporal variation. Additionally, because agriculture can increase environmental concentrations of Hg (Costantini, Czirják, Bustamante, Bumrungsri, & Voigt, 2019; Farella et al., 2007), we also tested whether more recently sampled bats within this rapidly changing landscape had higher Hg exposure. Second, using blood samples, we tested whether elevated bat Hg is correlated with impaired immune function. Expanding our species-specific analyses in vampire bats *(Desmodus rotundus;* Becker et al., 2017) across the entire bat community, we predicted that individuals with high Hg would have lower measures of cellular immunity. Third, we assessed infection with two bacterial pathogens, hemoplasmas and *Bartonella* spp. Both are common in Neotropical bats (Ikeda et al., 2017), including in these Belize sites (Becker, Bergner, et al., 2018; Becker, Speer, et al., 2020), and we previously showed infection can correlate with immunity in vampire bats specifically (Becker, Czirják, et al., 2018). However, how Hg shapes infection risk, and if such patterns are mediated by immunological relationships, is unknown. If Hg is associated with lower cellular immunity across bat species, we would expect greater concentrations to manifest in higher infection risks.

## Methods

### Bat sampling

During April 28 to May 4 2014 and April 24 to May 6 2017, we sampled 249 bats from 29 species captured in two areas of the Orange Walk District of Belize: Lamanai Archeological Reserve (LAR) and Ka’Kabish (KK). The LAR is bordered by the New River Lagoon, forest, and agriculture, while KK is a remnant forest patch surrounded by agriculture located 10 km away. At least 44 of the 70 bat species in Belize have been recorded in this region (Herrera, Duncan, Clare, Fenton, & Simmons, 2018; Reid, 1997). Bats were captured with mist nets from 19:00 until 22:00, and harp traps were also set from 18:00 to 05:00.

Bats were placed in individual cloth bags until processing and were identified to species based on morphology, including but not limited to body mass and forearm length (Reid, 1997). For Hg analysis, we trimmed <10 mg of fur from the dorsal or ventral region. Scissors were cleaned with ethanol between processed bats, and samples were stored in individual cryovials or Ziploc bags. From a subset of bats sampled in 2017, we collected 3–30 μL of blood by lancing the propatagial vein with sterile needles (23–30G; size and volume were dependent on bat mass), followed by collection using heparinized capillary tubes. Thin blood smears were prepared on glass slides and stained with Wright–Geimsa (Astral Diagnostics Quick III) to characterize cellular immunity. Remaining blood was stored on Whatman FTA cards (room temperature) or RNAlater (room temperature for four weeks and then –20°C) to preserve DNA. All bats were released after processing. Sampling followed guidelines for safe and humane handling of bats issued by the American Society of Mammalogists (Sikes, Care, & Mammalogists, 2016) and was approved by the University of Georgia Animal Care and Use Committee (A2014 04-016-Y3-A5). Sampling was authorized by the Belize Forest Department under permits CD/60/3/14(27), WL/2/1/17(16), and WL/2/1/17(19). Sample size for Hg varied by year (2014=98, 2017=148) and site (LAR=163, KK=83) and ranged from 1–58 individuals per species 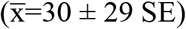.

### Fur Hg analysis

Bat fur was analyzed for total Hg (THg) at the Texas Christian University Aquatic Ecology Laboratory. THg data from 2014 were published previously (Becker, Chumchal, et al., 2018). Fur samples were rinsed in a 2:1 chloroform:methanol solution and dried overnight in a fume hood and reported on a fresh weight basis. We quantified THg with a direct Hg analyzer (DMA-80, Milestone, CT, USA) and analyzed National Research Council Canada reference material DORM 4 (certified value = 0.412 ± 0.036 mg/kg) every 10 samples for quality control; mean recovery was 94.32 ± 0.96% for 2017 data. Limited fur resulted in some samples falling below detection limit (DL), which was higher in 2014 (~0.48 ng). THg values below DL were estimated as 50% DL, and we used the 2014 DL to standardize concentrations (Rainwater et al., 2005). Fur THg was expressed in mg/kg and log_10_-transformed prior to statistical analyses. THg is a proxy for MeHg, which comprises 71-95% of Hg in bat fur (Yates et al., 2013).

### Statistical analysis of fur THg

To first link bats to aquatic food webs (i.e., the primary source of dietary Hg exposure; Becker, Chumchal, et al., 2018), we used the EltonTraits database to classify bat species according to the proportion of diet consisting of potentially aquatic prey: invertebrates, ectothermic tetrapods, and fish (Wilman et al., 2014). We used phylogenetic generalized linear mixed models (GLMMs) to test how THg varied with bat diet, site (LAR and KK), and year (2014 and 2017). We fit candidate models that considered all fixed effects and their two- and three-way interactions. We fit the phylogenetic GLMM using the *brms* package in R, default priors, and Gaussian errors (Bürkner, 2017). We included random effects for species and phylogeny, the latter of which used a phylogenetic covariance matrix derived from the Open Tree of Life through the *rotl* and *ape* packages (Michonneau, Brown, & Winter, 2016; Paradis, Claude, & Strimmer, 2004). We ran four chains for 20,000 iterations with a burn-in period of 10,000, thinned every 10 steps, for a total 4,000 samples. We compared GLMMs using leave-one-out cross-validation (LOOIC) and assessed fit with a Bayesian *R^2^*, including the total modeled variance and that attributed to only the fixed effects (Gelman, Goodrich, Gabry, & Vehtari, 2019; Vehtari, Gelman, & Gabry, 2017). We then estimated fixed effects per predictor (means and 95% highest density intervals [HDI]) and visualized fitted values using 100 random draws from the GLMM posterior distribution.

### Quantifying immunity and bacterial infection

Using blood smears, we first estimated total white blood cell (WBC) counts as the mean number of leukocytes from 10 fields of view (400X) with a light microscope (Schneeberger, Czirják, & Voigt, 2013). We next used differential WBC counts (1000X) to quantify the relative abundance of neutrophils and lymphocytes from 100 leukocytes. Neutrophils are components of the innate immune system, whereas lymphocytes are involved in adaptive responses like immunoglobulin production (Lanier, 2013). We derived absolute neutrophil and lymphocyte counts separately by multiplying total and differential WBCs. Elevated WBC counts can indicate a more robust cellular defense or an inflammatory response to acute infection.

To detect bacterial pathogens, we extracted genomic DNA from blood using QIAamp DNA Investigator Kits and DNeasy Blood and Tissue Kits (Qiagen). Before extracting RNAlater-preserved blood, samples were vortexed with 1 mL 1X phosphate-buffered saline and centrifuged for three minutes at 4000 RPM and 12 °C to prevent bacteria from floating in RNAlater; 200 μL from the bottom of the tube was then used for extraction. We used PCR to test for hemoplasmas (targeting the 16S rRNA gene) and *Bartonella* spp. (targeting the *gltA* gene) using previous diagnostic protocols (Bai, Gilbert, Fox, Osikowicz, & Kosoy, 2016; Volokhov et al., 2017). Hemoplasma data and sequences have been published previously (Becker, Speer, et al., 2020). Efforts to characterize *Bartonella* spp. in this bat community are ongoing, but prior studies of Belize vampire bats indicate high *gltA* sequence similarity to sequences from vampire bats in Mexico, Neotropical bat flies, and other Neotropical bats (Becker, Bergner, et al., 2018).

### Analyses of THg, immunity, and infection

We first used phylogenetic GLMMs to test the overall relationship between THg and both absolute neutrophil and lymphocyte counts. Each model included THg as the fixed effect, alongside sex and body condition (mass/forearm length) as covariates that could also affect leukocyte counts, with species and phylogeny as random effects. We used a Gaussian distribution for log_10_-transformed WBC counts. We next fit phylogenetic GLMMs with binomial errors to test associations between THg and infection with hemoplasmas and *Bartonella* spp. We then used causal mediation analyses (CMA) to test support for directional relationships among THg, immunity, and infection (Imai, Keele, & Tingley, 2010). Similar to structural equation modeling, CMA decomposes the hypothesized causal relationship between a predictor (i.e., THg) and a response (i.e., infection) into the direct effect and the indirect effect mediated through a third variable (i.e., immunity; Fig. S1). CMA then estimates the proportion of the total effect mediated through the indirect effect using a mediator model, which was each of the GLMMs predicting leukocyte counts by THg, and an outcome model. Here, we fit two GLMMs per pathogen that modeled infection as a function of THg and each WBC count. We then used the *brms* and *sjstats* packages to estimate these direct and indirect effects and in turn the proportion of the total relationship (THg and infection) mediated by the indirect relationship between THg and immunity and immunity and infection.

Because bat ecology likely dictates exposure to contaminants and pathogens, we assessed species-specific patterns in Hg-infection relationships. We fit logistic regressions for each bat species and pathogen when sample sizes were greater than two individuals. As these small samples can bias odds ratio estimates, we used the *logistf* package to implement Firth’s bias reduction (Heinze & Schemper, 2002). For species with no variance in infection, we assigned log odds of zero. Across these species, we next estimated phylogenetic signal (Pagel’s λ) in the log odds using the *caper* package and used phylogenetic generalized least squares to test if log odds covaried with sample size (Orme, 2013). We then used phylogenetic factorization to identify clades with different log odds at various taxonomic depths. We used the *taxize* and *phylofactor* packages to obtain taxonomic information for each species and partition log odds as a Gaussian response in a GLM (Chamberlain & Szöcs, 2013; Washburne et al., 2019). We included sample size as a weighting variable and used Holm’s sequentially rejective test to determine the number of significant clades.

## Results

### Neotropical bat fur THg

Expanding our initial studies of fur THg across this Neotropical bat community in 2014, we found that THg varied up to five orders of magnitude across species, sites, and years (Fig. 1). Our top GLMM included interactions between diet and year and between year and site (Table 1). Across years, the proportion of potentially aquatic prey in diet positively predicted THg but more so in 2014 (β=0.019, 95% HDI: 0.011-0.026) than in 2017 (β=0.016, 95% HDI: 0.008-0.023; Fig. 2A). We also identified strong spatiotemporal variation, such that fur THg increased between 2014 and 2017 for KK 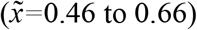 but slightly decreased for LAR 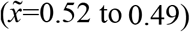.

**Figure 1.**
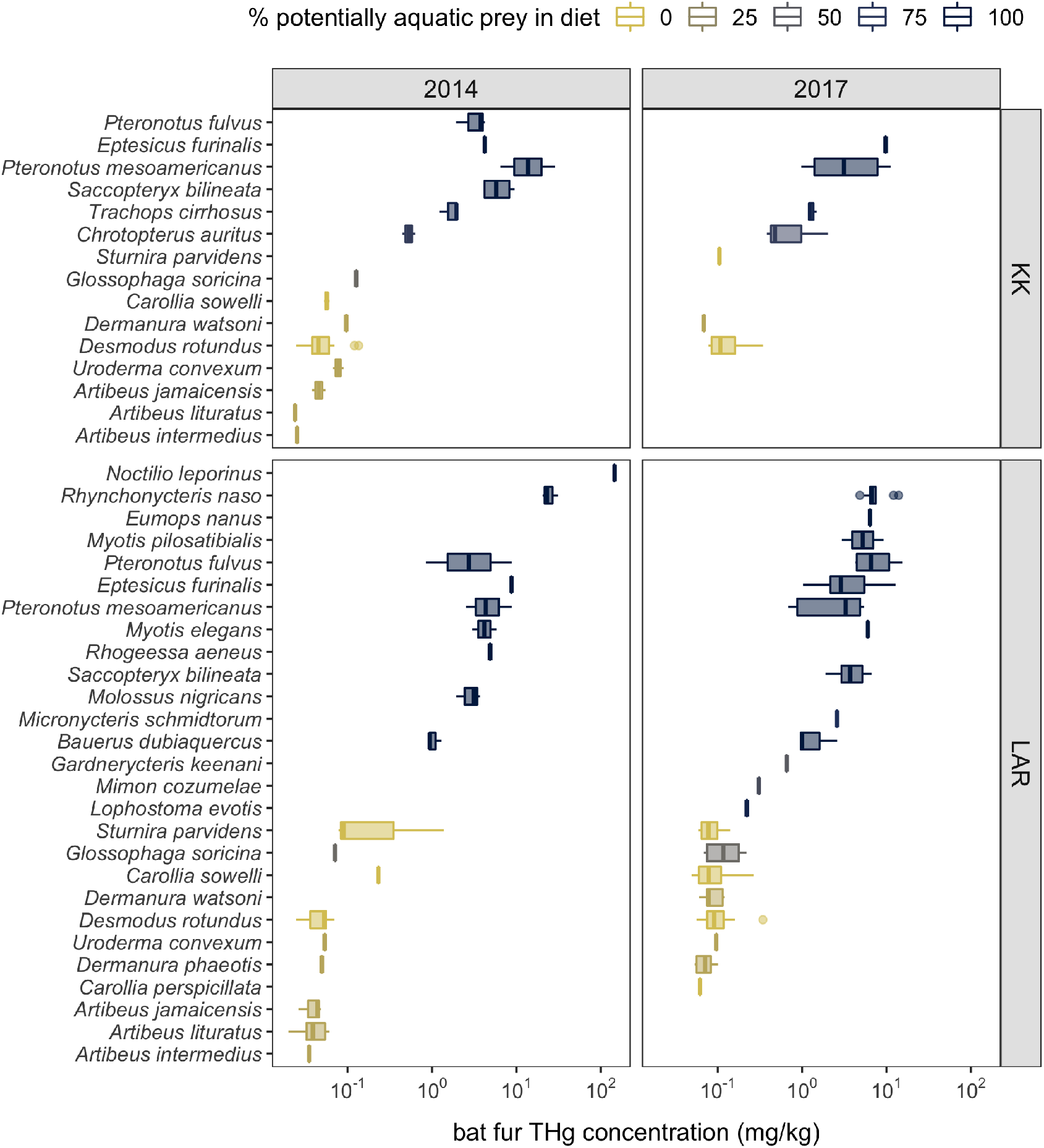
THg concentrations in bat fur across Neotropical bat species sampled in 2014 and 2017 across two sites in Belize. Boxplots are colored by the proportion of potentially aquatic prey in bat diets from the EltonTraits database. THg concentrations are displayed on a log_10_ scale.

**Figure 2.**
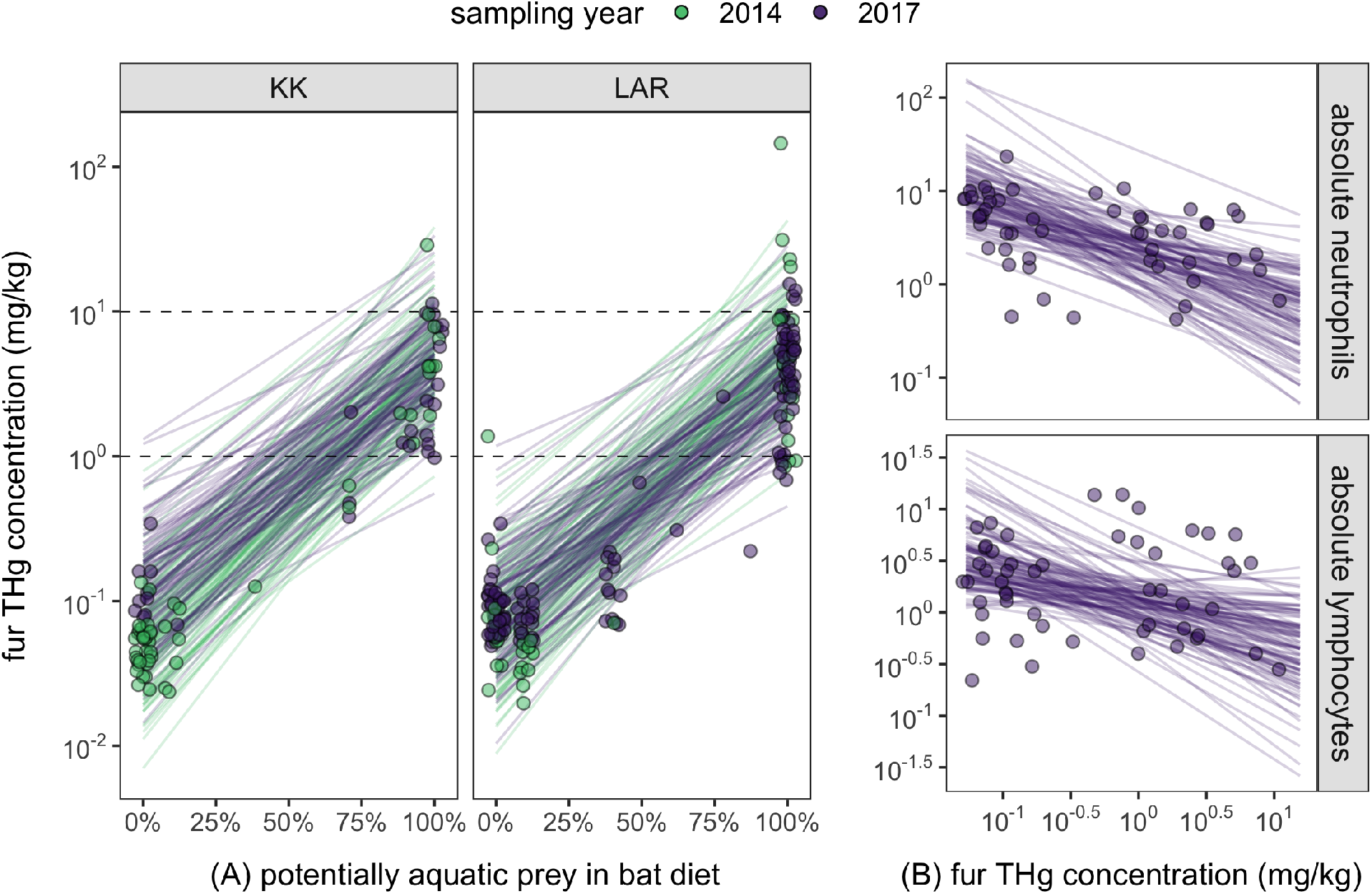
Dietary drivers of THg concentrations in Neotropical bat fur (A) and cellular immunity correlates of THg (B). Points indicate individual bats and are colored by sampling year in Belize. Lines display 100 random draws from the posterior distribution of the main phylogenetic GLMMs. THg concentrations and absolute leukocyte counts are displayed on a log_10_ scale.

**Table 1.**
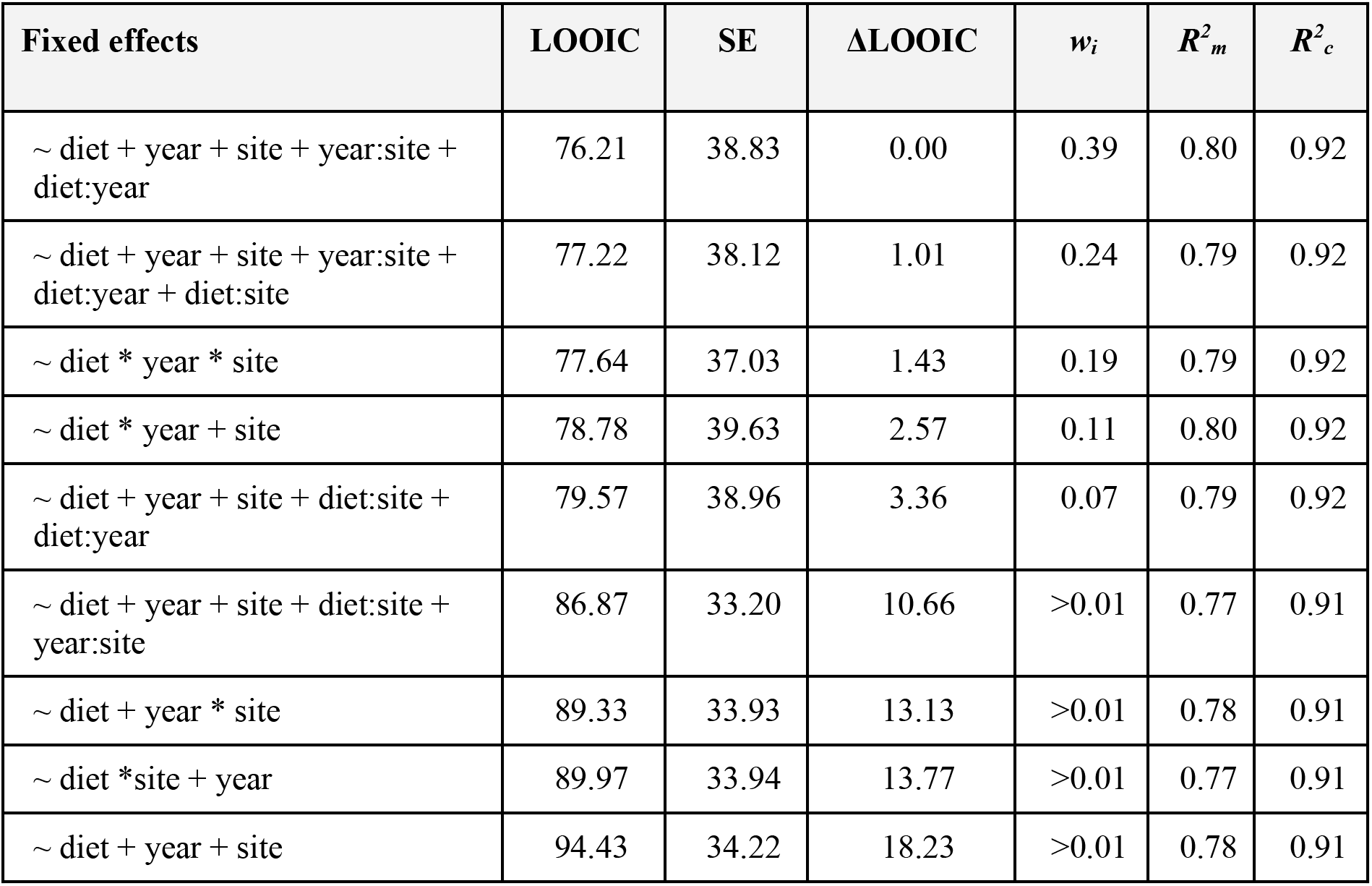
Competing phylogenetic GLMMs predicting log fur THg concentrations across the Belize bat community (*n*=249). Models are ranked by ΔLOOIC with LOOIC SE, Akaike weights (*w_i_*) and Bayesian *R^2^* estimates. All models include random effects for species and phylogeny.

### Associations with immunity and infection

Across bats sampled in 2017 with Hg and WBC data (*n*=52), our GLMMs showed that higher THg was associated with fewer neutrophils (β=–0.53, 95% HDI:–0.94 to –0.17) but not lymphocytes (β=–0.37, 95% HDI: –0.85 to 0.02; Fig. 2B) after adjusting for sex and condition; these two covariates had weak associations with leukocytes (Table S1). However, higher THg was also associated with lower odds of infection for hemoplasmas (OR=0.19, 95% HDI: 0.04– 0.79, *n*=134) and *Bartonella* spp. (OR=0.18, 95% HDI: 0.03–0.84, *n*=117; Fig. 3A). Accordingly, CMAs found that the relationship between THg and either WBC did not mediate any of the relationships between THg and infection for both pathogens (Table S2).

**Figure 3.**
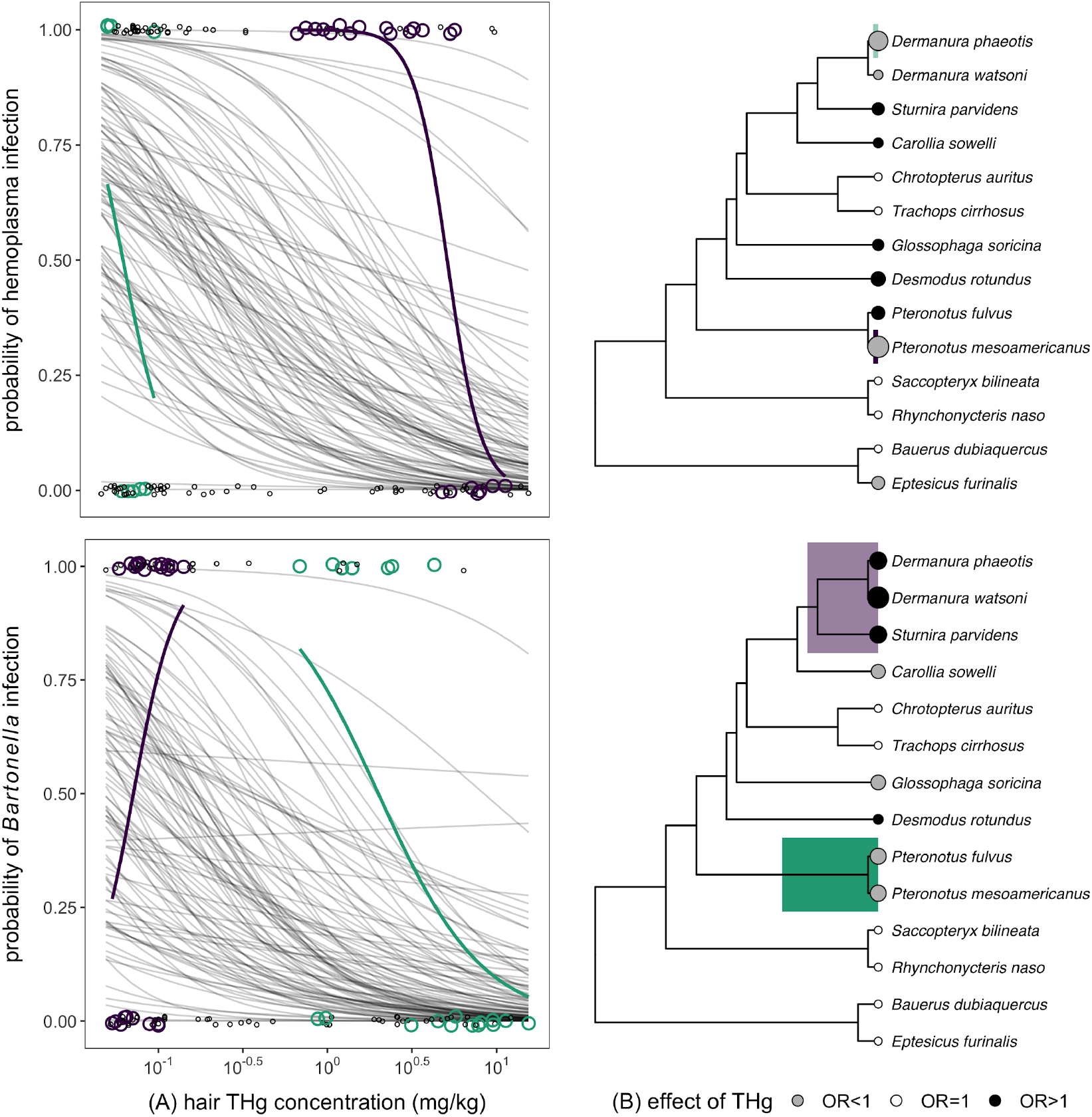
Associations between fur THg and bacterial infection (A) across the Belize Neotropical bat community and (B) on a per-species basis. Points in A indicate individual bats, and thin lines show 100 random draws from the posterior distribution of the phylogenetic GLMMs. Points in B indicate bat species, are scaled by the absolute log odds, and are colored by the direction of the relationship between THg and infection: null effects (i.e., OR=1, white), protective effects of THg (i.e., OR<1, grey), and mercury as a risk factor (i.e., OR>1, black). Data are shown for hemoplasmas (top) and *Bartonella* (bottom). Color indicates bat clades identified through phylogenetic factorization of log odds ratios, with lines indicating clade-specific GLM fits.

When we used logistic regression to analyze individual relationships between THg and infection per bat species and pathogen, we only found significant protective effects for *Pteronotus mesoamericanus* and hemoplasmas after adjusting for multiple comparisons (ln(OR)=–10.20, *p*=0.001; Table 2). We also detected a strong negative association between THg and *Bartonella* in this species (ln(OR)=–3.34), but no log odds were significantly different from zero after adjustment. Most species instead had null relationships between THg and infection (i.e., 36% for hemoplasmas and 43% for *Bartonella* spp.) or relatively weaker negative effects (e.g., *Dermanura phaeotis*, ln(OR)=–7.58 for hemoplasmas; *P. fulvus*, ln(OR)=–3.35 for *Bartonella* spp.). These negative THg–infection associations were identified for 29% of species for both pathogens. We also estimated similar proportions of positive THg associations for hemoplasmas (36%) and *Bartonella* spp. (29%). *Desmodus rotundus* had the strongest positive THg effect size for hemoplasma infection (ln(OR)=2.40), whereas *Dermanura watsoni* had the largest positive THg effect size for *Bartonella* spp. infection (ln(OR)=8.9).

**Table 2.** Results of species-specific logistic regressions (using Firth’s bias reduction method) between bat fur THg concentrations and infection status per each bacterial pathogen.

|  | Hemoplasmas |  |  |  | Bartonella |  |  |  |
| --- | --- | --- | --- | --- | --- | --- | --- | --- |
| Bat species | ln(OR) | *p* | $\sigma^2$ | *n* | ln(OR) | *p* | $\sigma^2$ | *n* |
| *Eptesicus furinalis* | -1.545 | 0.4569 | 0.214 | 8 | 0 | 1 | 0 | 7 |
| *Bauerus dubiaquercus* | 0 | 1 | 0 | 3 | 0 | 1 | 0 | 3 |
| *Dermanura watsoni* | -0.128 | 0.984 | 0.267 | 6 | 8.904 | 0.3177 | 0.167 | 6 |
| *Dermanura phaeotis* | -7.578 | 0.2562 | 0.278 | 9 | 4.27 | 0.5789 | 0.214 | 8 |
| *Sturnira parvidens* | 1.099 | 0.7923 | 0.257 | 15 | 4.198 | 0.4613 | 0.194 | 9 |
| *Carollia sowelli* | 0.213 | 0.9427 | 0.178 | 10 | -1.987 | 0.5159 | 0.233 | 10 |
| *Trachops cirrhosus* | 0 | 1 | 0 | 3 | 0 | 1 | 0 | 3 |
| *Chrotopterus auritus* | 0 | 1 | 0 | 3 | 0 | 1 | 0 | 3 |
| *Glossophaga soricina* | 0.665 | 0.8421 | 0.218 | 11 | -2.373 | 0.4911 | 0.194 | 9 |
| *Desmodus rotundus* | 2.395 | 0.3328 | 0.208 | 22 | 0.178 | 0.933 | 0.208 | 22 |
| *Pteronotus mesoamericanus* | -10.201 | 0.0001 | 0.227 | 22 | -3.339 | 0.0159 | 0.25 | 16 |
| *Pteronotus fulvus* | 1.653 | 0.65 | 0.25 | 4 | -3.346 | 0.4073 | 0.25 | 4 |
| *Rhynchonycteris naso* | 0 | 1 | 0 | 4 | 0 | 1 | 0 | 3 |
| *Saccopteryx bilineata* | 0 | 1 | 0 | 8 | 0 | 1 | 0 | 8 |

Comparative analyses of the log odds ratios across bat species revealed no phylogenetic signal for the relationship between THg and hemoplasmas (λ=0) but strong phylogenetic signal for the relationship between THg and *Bartonella* spp. (λ=0.84). Log odds ratios were not associated with sample size for hemoplasmas (*F*_1,12_=1.46, *p*=0.25) or *Bartonella* spp. (*F*_1,12_=0.25, *p*=0.63). Phylogenetic factorization further identified species-specific or taxonomic patterns in the magnitude and direction of effect size. For hemoplasmas, the odds of infection were significantly lower for *Pteronotus mesoamericanus* and *Dermanura phaeotis* when compared to the remaining sampled bat phylogeny. For *Bartonella* spp., however, the odds of infection were significantly lower for the genus *Pteronotus* (mean ln(OR)=-3.34) and significantly higher for the subfamily Stenodermatinae (*Dermanura* spp. and *Sturnira parvidens*, mean ln(OR)=5.79; Fig. 3B). Post-hoc GLMs showed that bats in the Stenodermatinae had especially negative associations between THg and neutrophils (β=−2.14, *p*=0.001) but not lymphocytes (β=−0.43, *p*=0.63), although these analyses were limited by small sample size (*n*=7).

## Discussion

Interactions among contaminants, immunity, and infection are difficult to disentangle in natural systems, but quantifying their proposed causal relationships can inform how land-use change affects wildlife health and human disease risks. By capitalizing on a diverse Neotropical bat system with high variation in Hg bioaccumulation and bacterial pathogens, we found higher THg was associated with fewer neutrophils but also lower odds of infection across the host community. However, our species-specific and taxonomic analyses showed THg had protective effects for hemoplasmas and *Bartonella* spp. in the genus *Pteronotus*, whereas THg was associated with fewer neutrophils and elevated infection in the subfamily Stenodermatinae. These contrasting relationships suggest contaminant-driven loss of pathogen habitat (i.e., anemia) or vector mortality versus immunosuppression as possible causal mechanisms, respectively, and identify clades of bats that may be especially resilient or vulnerable to infection risks from Hg exposure. Such findings more generally suggest contaminants may increase infection risk in some taxa but not others, emphasizing the importance of considering surveillance and management at different phylogenetic scales (Graham, Storch, & Machac, 2018).

Expanding our prior analyses of this Neotropical bat community and global patterns of THg in bat fur (Becker, Chumchal, et al., 2018) with larger within-species sample sizes, we first demonstrated that Hg exposure increases with potentially aquatic prey (or prey with some life stages linked to aquatic ecosystems) in diet despite spatial and temporal variation in THg. Positive associations with diets linked to aquatic ecosystems across sites and years provides additional support for trophic transfer of Hg through foraging (Cristol et al., 2008; Ortega-Rodriguez et al., 2019; Speir et al., 2014). This diet-mediated connectivity to aquatic ecosystems likely underlies other cases of guild-specific Hg bioaccumulation in bat communities (Carrasco-Rueda, Loiselle, & Frederick, 2020; Korstian, Chumchal, Bennett, & Hale, 2018). In many of these regions, bat dietary exposure to Hg is driven by land-use changes such as gold mining and agriculture (Carrasco-Rueda et al., 2020; Costantini et al., 2019), whereas atmospheric deposition is often the primary source of Hg in regions located further from anthropogenic point sources (Chételat et al., 2018; Korstian et al., 2018). The latter is a likely source of Hg in the Belize system (Becker, Chumchal, et al., 2018). However, intensified agriculture and especially slash-and-burn practices could provide other Hg inputs and may explain why bat THg increased between 2014 and 2017 in KK, the more agricultural site, but not in the protected LAR (Farella et al., 2007; Patterson, 2016).

Across bat species, fur THg was negatively correlated with neutrophil counts, which mirrors captive results and suggests impaired innate immunity (Lalancette, Morin, Measures, & Fournier, 2003). Previously, vampire bats sampled in Belize with high fur THg had weaker innate defense (i.e., bacterial killing ability; Becker et al., 2017). Wrinkle-lipped free-tailed bats with higher Hg exposure also had weaker innate immunity (i.e., bacterial killing ability, lysozyme and haptoglobin concentrations; Costantini et al., 2019). Most individual bats for which we had both Hg and immunity data showed THg below toxicity and subclinical thresholds of 5–10 mg/kg (Nam et al., 2012), which suggests innate immunity could be weakened at sublethal contaminant concentrations (Lewis, Cristol, Swaddle, Varian-Ramos, & Zwollo, 2013). Alternatively, sublethal effects of THg could combine with other stressors (e.g., reproduction, food and roost availability) to impair immunity. Additional immune measures across broader land-use gradients would help characterize functional differences in defense in relation to THg concentrations in anthropogenic habitats (Becker, Albery, et al., 2020; Costantini et al., 2019).

Although neutrophils were lowest in bats with high Hg exposure, the odds of infection with hemoplasmas and *Bartonella* spp. decreased with fur THg across the bat community. As in other mammals, bats challenged with bacteria produce neutrophils as part of the innate immune response (Weise, Czirják, Lindecke, Bumrungsri, & Voigt, 2017). Elevated neutrophil counts could be protective, as implied through negative associations between innate immunity and these two pathogens in vampire bats (Becker, Czirják, et al., 2018). Accordingly, relationships between THg and immunity did not mediate the relationships between THg and infection, which likely indicates immunosuppression is not a causal mechanism. One alternative may involve contaminant-mediated pathogen mortality; for example, lead exposure likely caused helminth mortality in avian hosts (Prüter et al., 2018). As facultative intracellular pathogens, hemoplasmas and *Bartonella* spp. both infect red blood cells. Hg can lower erythrocyte counts (Shaw, Dash, & Panigrahi, 1991), which could reduce resources available to both bacteria. We did not measure anemia or Hg in blood; however, fur THg strongly correlates with blood THg, despite the former being orders of magnitude higher than the latter (Wada, Yates, Evers, Taylor, & Hopkins, 2010). As *Bartonella* spp. and possibly hemoplasmas likely depend in part on vector-borne transmission (Becker, Bergner, et al., 2018; Willi et al., 2007), ectoparasite mortality from contaminants in hosts or the environment could also explain observed infection patterns (e.g., as found for some avian ectoparasites; Eeva & Klemola, 2013). Lastly, we did not reliably age all bats, precluding age from our analyses; however older animals typically have higher fur THg (Korstian et al., 2018). If older bats also have stronger adaptive immunity (e.g., antibody titers in *Saccopteryx bilineata*; Schneeberger, Courtiol, Czirják, & Voigt, 2014), age could explain negative associations between THg and infection.

Our species-specific analyses suggested contaminants may have variable impacts on infection risk depending on taxon. In particular, we identified possibly protective effects of THg on infections primarily in the genus *Pteronotus*, with strong negative effects for *P. mesoamericanus* for both pathogens. Both species in this genus had high THg, being insectivores that often eat dipterans and coleopterans (Salinas-Ramos, Herrera Montalvo, León-Regagnon, Arrizabalaga-Escudero, & Clare, 2015). Given this clade-based signal, *Pteronotus* bats may be particularly resilient to disease impacts of Hg. Such effects could be in part mediated by high host-specificity of bat flies for this genus (Ter Hofstede, Fenton, & Whitaker, 2004), particularly if high THg increases vector mortality or if aspects of bat fly biology impact vector competence (Weiss & Aksoy, 2011). In contrast, we also identified taxa with positive relationships between THg and infection. For hemoplasmas, higher THg was associated with greater risk in vampire bats. As this species frequently feeds on humans and livestock (Streicker & Allgeier, 2016), pathogen surveillance would be particularly important in habitats contaminated by agriculture or mining. Additionally, the Stenodermatinae displayed strong positive associations between THg and *Bartonella* spp. These frugivores also showed negative correlations between THg and neutrophils, suggesting Hg-mediated immunosuppression. Such species may thus be especially vulnerable to infection from Hg exposure and could play key roles in maintaining *Bartonella* spp. infection cycles between bats and ectoparasites in contaminated environments (Judson, Frank, & Hadly, 2015). From another perspective, because the Stenodermatinae and *Pteronotus* had lower and higher fur THg, respectively, low Hg exposure could increase susceptibility while higher concentrations instead cause anemia and facilitate protective effects against erythrocytic pathogens. More generally, these results highlight the importance of considering surveillance and management of Hg exposure (e.g., through possible land use drivers) at different phylogenetic scales, such as species, genus, or subfamily.

Beyond bats and their pathogens, our study more broadly emphasizes the need to assess potential causal relationships between contaminants and infectious diseases in natural systems. In particular, we provide a novel perspective on integrating approaches from ecotoxicology with those of disease ecology to disentangle the complex relationships among contaminants, immunity, and infection. Future work that carefully integrates data on contaminant exposure, specific diet composition, multiple immune measures, and pathogen diversity (e.g., with metagenomics; Bergner et al., 2019) across site gradients of anthropogenic intensity could help identify habitats, host clades, and infections for which disease risks are highest.

## Acknowledgements

For assistance with field logistics and permits, we thank Mark Howells, Neil Duncan, and staff of the Lamanai Field Research Center. We also thank the many colleagues who helped net bats during 2014 and 2017 bat research in Belize as well as Susan Perkins for laboratory reagents. Lastly, we thank two anonymous reviewers for constructive feedback on this manuscript.

## Data accessibility

Data from 2017 will be deposited in Dryad Digital Repository upon acceptance. Data from 2014 have been published previously (Becker, Chumchal, et al., 2018).

## Author contributions

DJB, KAS, AMB, HGB, and NBS collected samples; JMK, DJB, KAS, DVV, AMB, CLB, HD, and TPS analyzed samples; DJB analyzed the data; and HGB, RKP, TRR, MBF, NBS, and MMC provided funding and logistical support. DJB wrote the manuscript with input from all authors.

