## Supplemental Material for "Disentangling interactions among mercury, immunity, and infection in a Neotropical bat community"

Figure S1. Hypothesized causal pathway between fur THg concentrations, absolute WBC counts (neutrophils and lymphocytes), and bacterial infection status (hemoplasmas and *Bartonella*). CMA estimates the total indirect relationship between THg and immunity (*a*) and immunity and infection (*b*) as well as the direct relationship between THg and infection (*c'*). The total effect (*c*) is the sum of the indirect effect (*ab*) and the direct effect (*c'*). The proportion mediated by immunity ($P_M$) is derived as the indirect effect (*ab*) divided by the estimated total effect (*c*): *ab/c*.

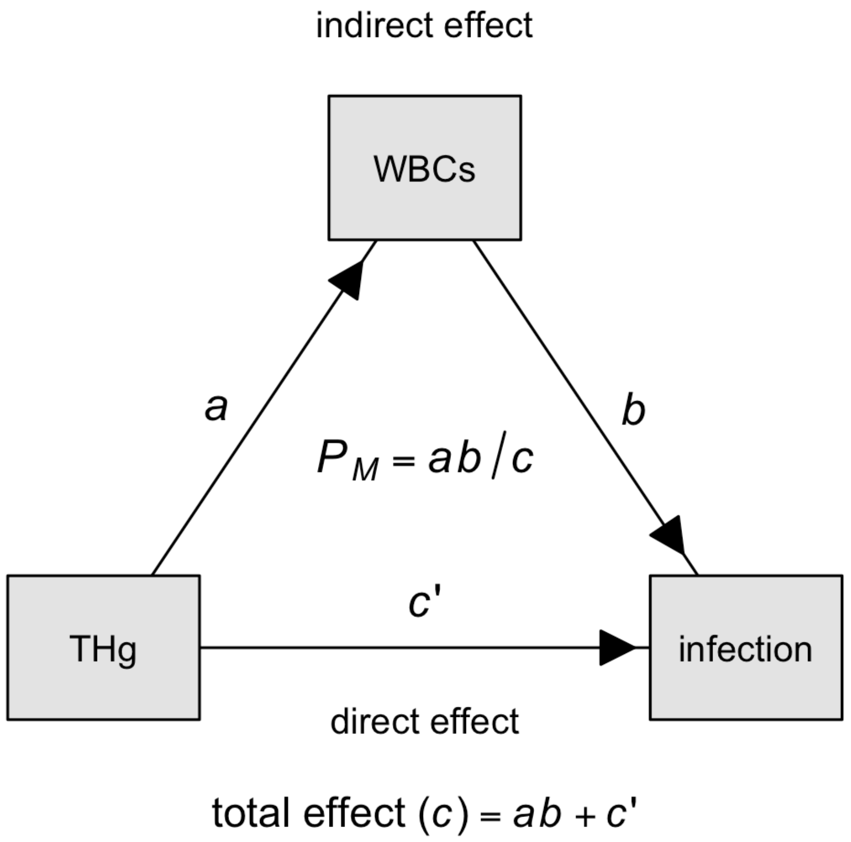

Table S1. Full results of the phylogenetic GLMMs predicting absolute leukocyte counts as a function of log fur THg concentration, bat sex, and bat body condition. All models include random effects for species and phylogeny. We report means and 95% HDIs for the estimated fixed effects, and estimates with 95% HDIs that do not cross zero are displayed in bold.

| | Neutrophils | | Lymphocytes | |
|---|---|---|---|---|
| Coefficient | Mean | 95% HDI | Mean | 95% HDI |
| Intercept | 0.54 | –0.25 to 1.26 | 0.07 | –0.73 to 0.78 |
| Log THg | **–0.53** | **–0.94 to –0.17** | –0.37 | –0.85 to 0.02 |
| Males | –0.15 | –0.41 to 0.10 | –0.11 | –0.44 to 0.22 |
| Condition | –0.14 | –1.20 to 0.88 | 0.26 | –0.87 to 1.44 |

Table S2. Results of the CMA among log fur THg, absolute leukocyte counts, and infection status. As described in Figure S1, we report the estimated direct (*c'*) and total indirect (*ab*) effect alongside the proportion mediated by leukocyte counts ($P_M$). All analyses used phylogenetic GLMMs with random effects for species and phylogeny using the *brms* and *sjstats* packages. Results are given as the posterior mean and 95% HDI for analyses considering neutrophil or lymphocyte counts as the mediator variable and with (*i*) hemoplasmas or (*ii*) *Bartonella* as the outcome variable. Estimates with 95% HDIs that do not cross zero are displayed in bold.

| | | Neutrophils as mediator | | Lymphocytes as mediator | |
|---|---|---|---|---|---|
| | Estimate | Mean | 95% HDI | Mean | 95% HDI |
| *i* | Direct effect | **–3.52** | **–7.02 to –0.60** | –2.29 | –5.13 to 0.10 |
| | Indirect effect | 0.43 | –0.55 to 1.89 | –0.38 | –1.52 to 0.21 |
| | Proportion mediated | –0.14 | –0.81 to 0.53 | 0.14 | –0.27 to 0.54 |
| *ii* | Direct effect | **–4.31** | **–8.73 to –0.82** | **–3.24** | **–6.80 to –0.35** |
| | Indirect effect | 0.76 | –0.26 to 2.57 | 0.16 | –0.37 to 1.31 |
| | Proportion mediated | –0.22 | –1.04 to 0.60 | –0.05 | –0.50 to 0.40 |
